# SARS-CoV-2 Whole Genome Amplification and Sequencing for Effective Population-Based Surveillance and Control of Viral Transmission

**DOI:** 10.1101/2020.06.06.138339

**Authors:** Divinlal Harilal, Sathishkumar Ramaswamy, Tom Loney, Hanan Al Suwaidi, Hamda Khansaheb, Abdulmajeed Alkhaja, Rupa Varghese, Zulfa Deesi, Norbert Nowotny, Alawi Alsheikh-Ali, Ahmad Abou Tayoun

**Affiliations:** Al Jalila Genomics Center, Al Jalila Children’s Hospital, Dubai, United Arab Emirates; College of Medicine, Mohammed Bin Rashid University of Medicine and Health Sciences, Dubai, United Arab Emirates; Medical Education & Research Department, Dubai Health Authority, Dubai, United Arab Emirates; Microbiology and Infection Control Unit, Pathology and Genetics Department, Latifa Women and Children Hospital, Dubai Health Authority, Dubai, United Arab Emirates; Institute of Virology, University of Veterinary Medicine Vienna, Vienna, Austria

## Abstract

**Background:** With the gradual reopening of economies and resumption of social life, robust surveillance mechanisms should be implemented to control the ongoing COVID-19 pandemic. Unlike RT-qPCR, SARS-<u>C</u>oV-2 <u>W</u>hole <u>G</u>enome <u>S</u>equencing (cWGS) has the added advantage of identifying cryptic origins of the virus, and the extent of community-based transmissions versus new viral introductions, which can in turn influence public health policy decisions. However, practical and cost considerations of cWGS should be addressed before it can be widely implemented.

**Methods:** We performed shotgun transcriptome sequencing using RNA extracted from nasopharyngeal swabs of patients with COVID-19, and compared it to targeted SARS-CoV-2 full genome amplification and sequencing with respect to virus detection, scalability, and cost-effectiveness. To track virus origin, we used open-source multiple sequence alignment and phylogenetic tools to compare the assembled SARS-CoV-2 genomes to publicly available sequences.

**Results:** We show a significant improvement in whole genome sequencing data quality and viral detection using amplicon-based target enrichment of SARS-CoV-2. With enrichment, more than 99% of the sequencing reads mapped to the viral genome compared to an average of 0.63% without enrichment. Consequently, a dramatic increase in genome coverage was obtained using significantly less sequencing data, enabling higher scalability and significant cost reductions. We also demonstrate how SARS-CoV-2 genome sequences can be used to determine their possible origin through phylogenetic analysis including other viral strains.

**Conclusions:** SARS-CoV-2 whole genome sequencing is a practical, cost-effective, and powerful approach for population-based surveillance and control of viral transmission in the next phase of the COVID-19 pandemic.

---

The COVID-19 pandemic continues to inflict devastating human life losses (1), and has enforced significant social changes and global economic shut downs (2). With the accumulating financial burdens and unemployment rates, several governments are sketching out plans for slowly re-opening the economy and reviving social life and economic activity. However, robust population-based surveillance systems are essential to track viral transmission during the re-opening process.

While RT-qPCR targeting SARS-CoV-2 can be effective in identifying infected individuals for isolation and contact tracing, it is not useful in determining which viral strains are circulating in the community: autochthonous versus imported ones, and – if imported – it is important to know the origin of the strains, which in turn influences public health policy decisions. In addition, it is vital to identify super-spreader events as they can be influenced by the virus strain (3). SARS-CoV-2 whole genome sequencing (cWGS), on the other hand, can detect the virus and can delineate its origins through phylogenetic analysis (4, 5) in combination with other local and international viral strains, especially given the accumulation of thousands of viral sequences from countries all over the world (www.nextstrain.org) (**Figure 1**). However, practical considerations, such as cost, scalability, and data storage, should first be investigated to assess the feasibility of implementing cWGS as a population-based surveillance tool. Here we show that cWGS is cost-effective and is highly scalable when using a target enrichment sequencing method, and we also demonstrate its utility in tracking the origin of SARS-CoV-2 transmission.

**Figure 1.**
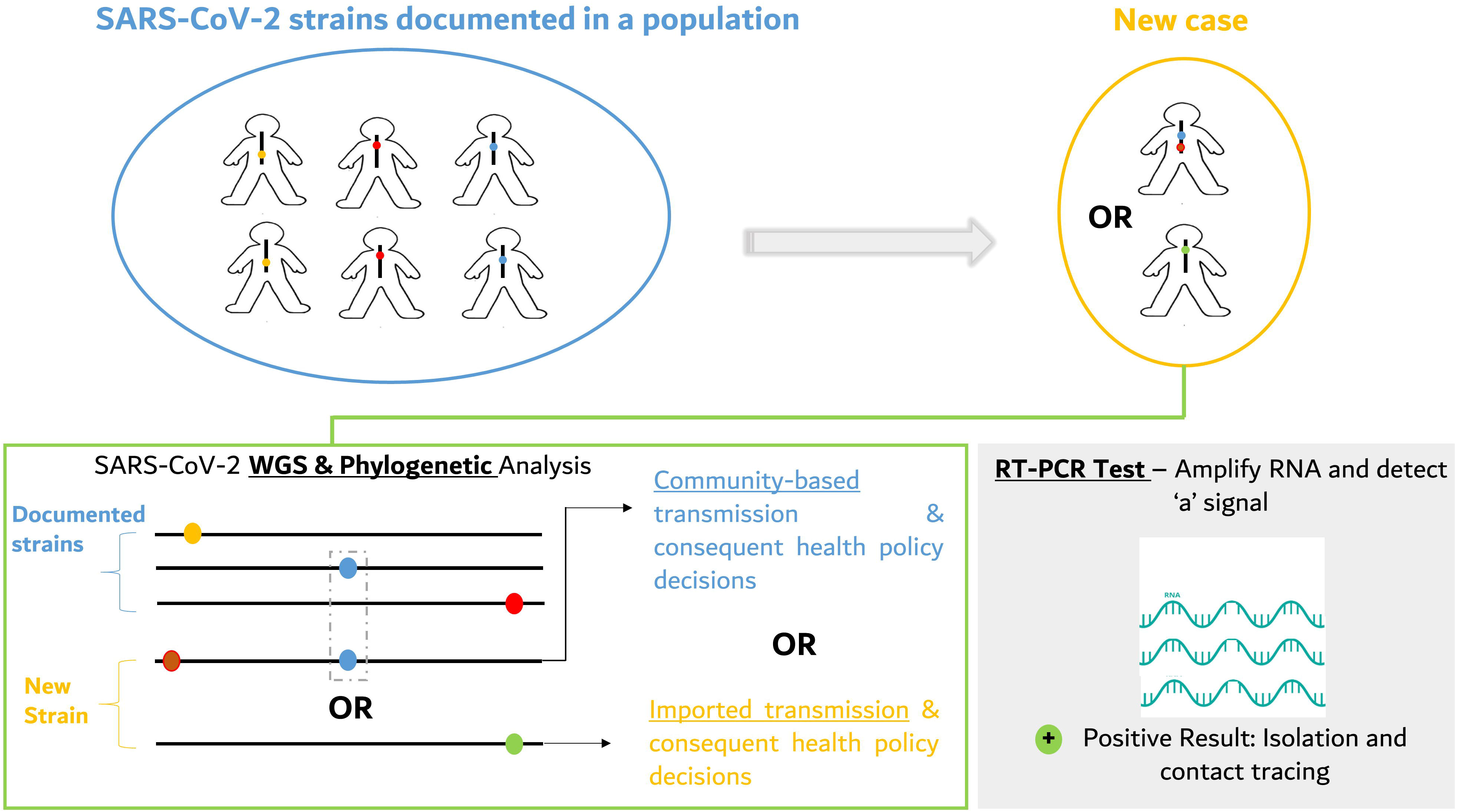
SARS-CoV-2 whole genome sequencing-based surveillance. A schematic illustrating how SARS-CoV-2 whole genome sequencing (cWGS) can be used as a surveillance tool to uncover community-based versus international/travel-related introductions. Mutations are represented by coloured dots or circles on SARS-CoV-2 genomes (black bars) within each patient with COVID-19. A population of viral genomes in a community can be used as a reference set (circled blue) for future analysis when new cases (circled orange) emerge. Two scenarios are represented for the new case: the first represents community transmission while the second represents external introduction. The strain representing community transmission has two mutations, one of which (blue) has been identified in a strain from a previous patient in this community, while the second is a new mutation (brick red), arising as part of the virus evolution. The strain with a single novel mutation (green) not seen previously in this population represents a new introduction.

## Materials and Methods

### Human subjects and ethics approval

All patients had laboratory-confirmed COVID-19 based on positive RT-PCR assay for SARS-CoV-2 in the Centralized Dubai Health Authority (DHA) virology laboratory. This study was approved by the Dubai Scientific Research Ethics Committee - Dubai Health Authority (approval number #DSREC-04/2020_02).

### RNA extraction and SARS-CoV-2 detection

Viral RNA was extracted from nasopharyngeal swabs of patients with COVID-19 using the EZ1 DSP Virus Kit (Qiagen, Hilden, Germany), optimized for viral and bacterial nucleic acids extractions from human specimens using magnetic bead technology. SARS-CoV-2 positive results were confirmed using a RT-PCR assay, originally designed by the US Centres for Disease Control and Prevention (CDC), and is currently provided by Integrated DNA Technologies (IDT, IA, USA). This assay consists of oligonucleotide primers and dual-labelled hydrolysis (TaqMan®) probes (5’FAM/3’Black Hole Quencher) specific for two regions (N1 and N2) of the virus nucleocapsid (N) gene. An additional primer/probe set is also included to detect the human RNase P gene (RP) as an extraction control. The reverse transcription and amplification steps are performed using the TaqPath™ 1-Step RT-qPCR Master Mix (ThermoFisher, MA, USA) following manufacturer’s instructions. A sample was considered positive if the cycle threshold (Ct) values were less than 40 for each of the SARS-CoV-2 targets (N1 and N2) and the extraction control (RP). To estimate the viral load relative to human RNA, we calculated the ΔCt value for each target as follows: ΔCt = Ct_Nn_ – Ct_RP_, where N_n_ is either N1 or N2. The average of the N1 and N2 target ΔCt values was then negated to reach a relative estimate of viral load which is inversely correlated with Ct value.

### Shotgun transcriptome SARS-CoV-2 sequencing

RNA libraries from all samples were prepared for shotgun transcriptomic sequencing using the TruSeq Stranded Total RNA Library kit from Illumina (San Diego, CA, USA), following manufacturer’s instructions. RNA specific fluorescent dye is used to quantify extracted RNA using the Qubit RNA XR assay kit and the Qubit Flourometer system (ThermoFisher, MA, USA). Then, 1μg of input RNA from each patient sample was depleted for human ribosomal RNA, and the remaining RNA underwent fragmentation, reverse transcription (using the SuperScript II Reverse Transcriptase Kit from Invitrogen, Carisbad, USA), adaptor ligation, and amplification. Libraries were then sequenced using the NovaSeq SP Reagent kit (2 × 150 cycles) from Illumina (San Diego, CA, USA).

### Targeted amplification and sequencing of SARS-CoV-2 genome

RNA extracted (~1μg) from patient nasopharyngeal swabs was used for double stranded cDNA synthesis using the QuantiTect Reverse Transcription Kit (Qiagen, Hilden, Germany) according to manufacturer’s protocol. This cDNA was then evenly distributed into 26 PCR reactions for SARS-CoV-2 whole genome amplification using 26 overlapping primer sets covering most of its genome (**Figure 2A** and **Supplemental Table 1**). The SARS-CoV-2 primer sets used in this study were modified from Wu *et al* (6) by adding M13 tails to enable sequencing by Sanger, if needed (**Supplemental Table 1**). PCR amplification was performed using the Platinum™ SuperFi™ PCR Master Mix (ThermoFisher, MA, USA) and a thermal protocol consisting of an initial denaturation at 98°C for 60 seconds, followed by 27 cycles of denaturation (98°C for 17 seconds), annealing (57°C for 20 seconds), and extension (72°C for 1 minute and 53 seconds). A final extension at 72°C for 10 minutes was applied before retrieving the final PCR products. Amplification was confirmed by running 2μl from each reaction on a 2% agarose gel.

**Figure 2.**
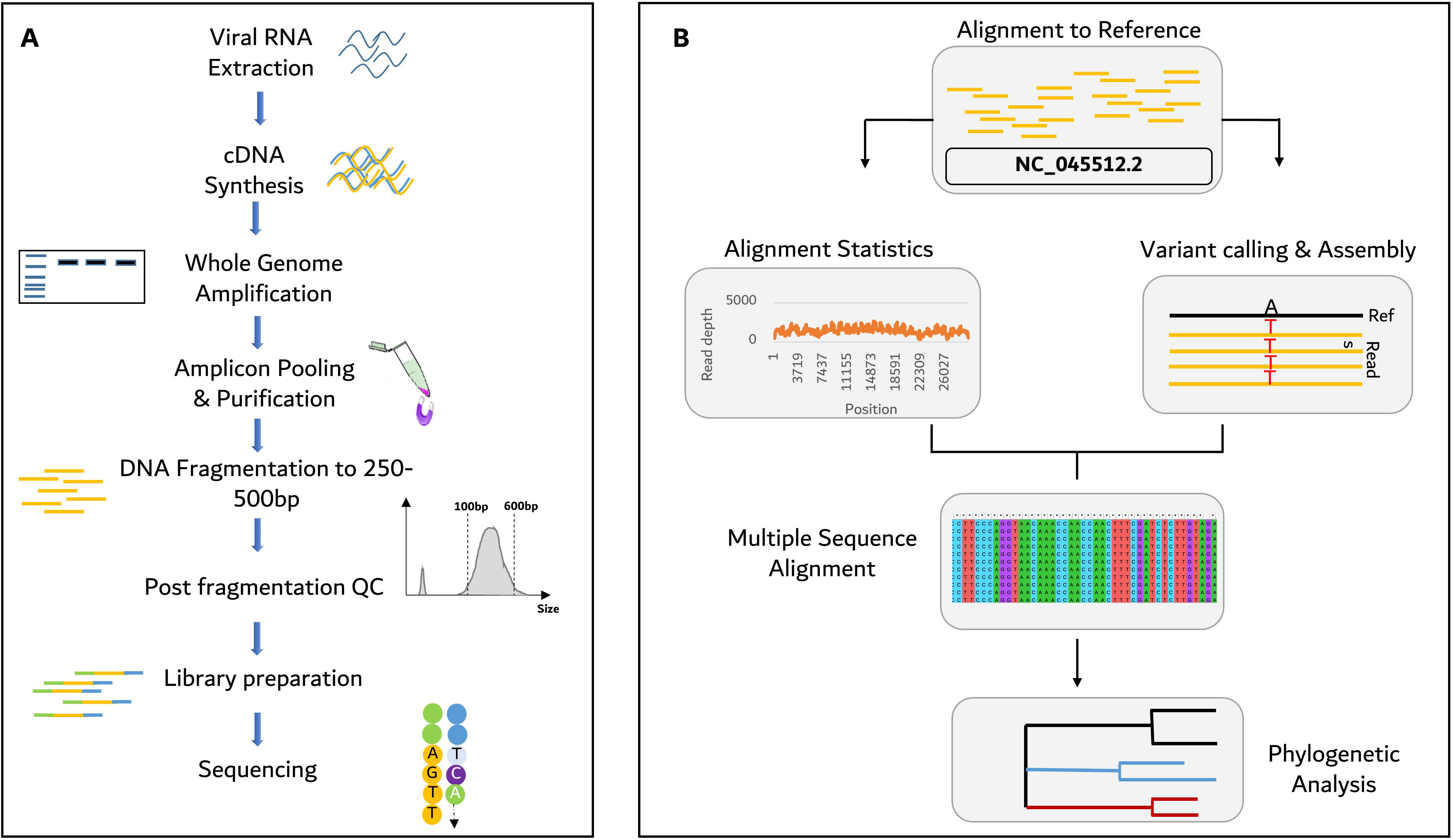
Whole genome amplification, sequencing, and phylogenetic analysis of SARS-CoV-2 genome. >A, Wet bench steps describing SARS-CoV-2 genome enrichment and sequencing. B, Bioinformatics and computational steps for sequence alignment, variant calling, SARS-CoV-2 genome assembly, multiple sequence alignment and phylogenetic analysis. All steps are described in details in Methods.

All PCR products were then purified using Agencourt AMPure XP beads (Beckman Coulter, CA, USA), quantified by NanoDrop (ThermoFisher, MA, USA), diluted to the same concentration, and then pooled into one tube for next steps.

A minimum of 200-800ng of the pooled PCR products in 55μl were then sheared by ultra-sonication (Covaris LE220-plus series, MA, USA) to generate a target fragment size of 250-750bp using the following parameters: 20% Duty Factor, Peak Power of 150 Watts, 900 cycles per burst, 320 seconds Treatment Time, an Average Power of 30 Watts, and 20°C bath temperature. Target fragmentation were confirmed by the TapeStation automated electrophoresis system TapeStation (Agilent, CA, USA) (**Figure 2A**). Subsequently, the fragmented product is purified and then processed to generate sequencing-ready libraries using the SureSelectXT Library Preparation kit (Agilent, CA, USA) following manufacturer instructions. Indexed libraries from multiple patients were pooled and sequenced (2 × 150 cycles) using the MiSeq or the NovaSeq systems (Illumina, San Diego, CA, USA). A step-by-step SARS-CoV-2 target enrichment and sequencing protocol is provided in **Appendix I**.

### Bioinformatics analysis and SARS-CoV-2 genome assembly

Demultiplexed Fastq reads, obtained through shotgun or target enrichment sequencing, were generated from raw sequencing base call files using BCL2Fastq v2.20.0, and then mapped to the reference Wuhan genome (GenBank accession number: NC_045512.2) by Burrow-Wheeler Aligner, BWA v0.7.17. Alignment statistics, such as coverage and mapped reads, were generated using Picard 2.18.17. Variant calling was performed by GATK v3.8-1-0, and was followed by SARS-CoV-2 genome assembly using BCFtools v.1.3.1 (**Figure 2B**).

All tools used in this study are freely accessible. For laboratories without bioinformatics support, several publicly accessible, end-to-end bioinformatics pipelines (INSaFlu: https://insaflu.insa.pt/; Genome Detective: https://www.genomedetective.com/app/typingtool/virus/) (7,8), composed of the above tools, can be used to generate viral sequences from raw Fastq data.

For downstream analysis, a general quality control metric was implemented to ensure assembled SARS-CoV-2 genomes have at least 20X average coverage (sequencing reads>Q30) across most nucleotide positions (56-29,797).

For target enrichment and shotgun sequencing comparisons in **Table 1**, we used data from 7 samples (UAE/P1/2020, L0287, L1189, L4711, L5857, L6841, L9119) generated using both methods. Data from patient L5630 were not included in this analysis since, unlike the above 7 samples, significantly more sequencing data was allocated for the target enriched sample in this patient (**Supplemental Table 2**) which can overestimate the efficiency of the target enrichment protocol.

**Table 1.** RT-PCR and transcriptome sequencing statistics for COVID-19 patients

| Sample ID | RT-PCR | Shotgun Transcriptome Sequencing data |  |  |  |  |
| --- | --- | --- | --- | --- | --- | --- |
|  | Viral RNA Abundance (-ΔCt) | Data size (Gb) | Total Reads | Reads Aligned* | % Reads aligned* | Mean Coverage* |
| L8205** | -0.995 | 4.80 | 32,117,004 | 124,373 | 0.38 | 10.13 |
| L4280 | 3.852 | 5.10 | 33,921,438 | 69,460 | 0.20 | 20.94 |
| L0826** | -12.065 | 3.21 | 21,383,276 | 57,143 | 0.26 | 2.77 |
| L2771** | -6.196 | 5.70 | 38,114,908 | 131,449 | 0.34 | 5.27 |
| L9440 | 3.258 | 4.20 | 28,015,394 | 107,206 | 0.38 | 38.39 |
| L1758 | 10.741 | 4.90 | 32,649,592 | 937,403 | 2.87 | 2106.37 |
| L0000 | 2.823 | 5.05 | 33,700,572 | 208,432 | 0.62 | 43.23 |
| L3779 | 5.345 | 4.49 | 29,912,422 | 395,560 | 1.32 | 320.98 |
| L5630** | -1.648 | 4.57 | 30,462,036 | 70,058 | 0.23 | 4.16 |
| L4184 | 1.635 | 3.77 | 25,150,950 | 215,797 | 0.86 | 31.14 |
| L0287** | 9.096 | 5.40 | 35,991,646 | 129,076 | 0.36 | 30.07 |
| L1189** | 7.344 | 5.70 | 37,968,228 | 72,599 | 0.19 | 7.85 |
| L4711** | 8.307 | 4.91 | 32,762,088 | 140,388 | 0.43 | 13.63 |
| L5857** | 10.824 | 3.86 | 25,757,828 | 162,345 | 0.63 | 49.44 |
| L6841** | 7.571 | 4.96 | 33,051,790 | 264,298 | 0.80 | 12.85 |
| L9119** | 9.221 | 5.36 | 35,719,244 | 97,709 | 0.27 | 19.10 |
| UAE/P1/2020** | 3.628 | 4.13 | 27,542,314 | 126,756 | 0.46 | 227.00 |
| Average | 3.691 | 4.71 | 31,667,401 | 198,956 | 0.63 | 173.14 |

\*Statistics with respect to the SARS-CoV-2 genome
\*\*Sequencing Data generated in this study. For the remaining samples, data was generated in Abou Tayoun et al study

All new SARS-CoV-2 sequences (n=7) generated in this study were submitted to GISAID (Global Initiative on Sharing All Influenza Data) under accession IDs: EPI_ISL_463740 and EPI_ISL_469276 to EPI_ISL_469281.

### Phylogenetic analysis

We used Nexstrain (9), which consists of Augur v6.4.3 pipeline for multiple sequence alignment (*MAFFT* v7.455) (10) and phylogenetic tree construction (*IQtree* v1.6.12) (11). Tree topology was assessed using the fast bootstrapping function with 1,000 replicates. Tree visualization and annotations were performed in *FigTree* v1.4.4 (12).

## Results

### SARS-CoV-2 whole genome sequencing

Shotgun transcriptome sequencing was used to fully sequence SARS-CoV-2 RNA extracted from patients (n=17) who tested positive for the virus (4). Analysis of the sequencing data showed that this approach required, on average, 4.71Gb of data per sample yielding 31.7 million total reads, of which approximately 0.63% of the reads (~199,000 reads) mapped to the SARS-CoV-2 genome with an average coverage of ~173x (**Table 1**). This is attributed to the fact that most of the shotgun data (~99%) is allocated to the human transcriptome while a minority of the reads align to the SARS-CoV-2 genome (**Table 1**). In addition to cost and storage considerations discussed below, this approach is not highly sensitive for detecting SARS-CoV-2 genomes in general and specifically in samples with low viral abundance. In fact, despite high viral abundance relative to human RNA, most samples had less than 100x sequencing coverage across the SARS-CoV-2 genome. Samples with seemingly very low viral loads failed to yield full SARS-CoV-2 genome sequence using this approach (**Table 1** and **Figure 3A**).

**Figure 3.**
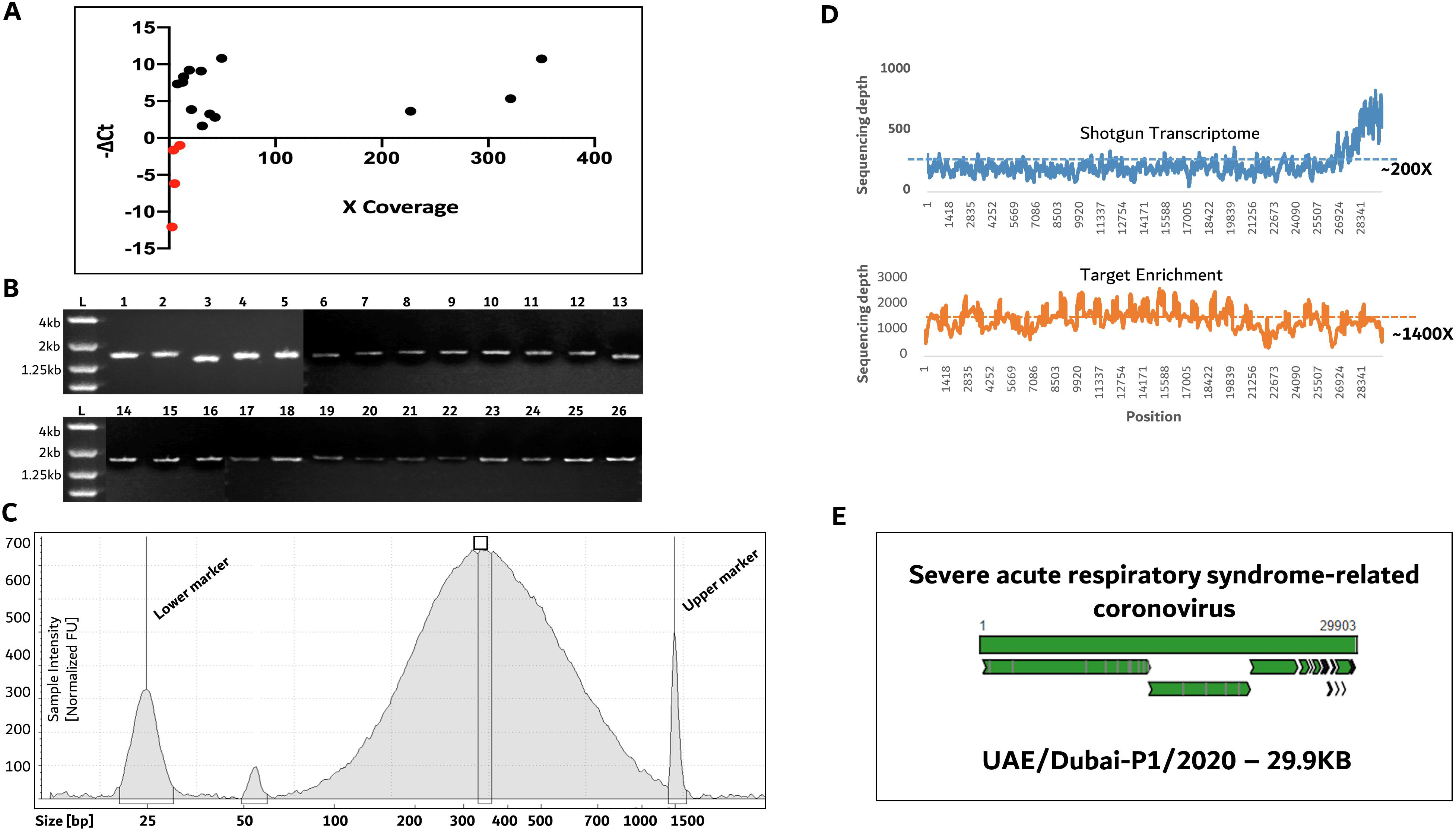
SARS-CoV-2 RNA detection, targeted enrichment, and full sequencing. A, Relationship between the RT-PCR cycling threshold and sequencing coverage over the SARS-CoV-2 genome. –ΔCt is calculated as an estimate of viral load relative to human RNA (see Methods). Red circles represent lowest –ΔCt values (and lowest relative viral abundance) from samples with very low sequencing coverage. Sequencing data were generated by the shotgun method. B, an agarose gel showing the overlapping 26 PCR products (~1.5kb) covering the SARS-CoV-2 genome. C, An electrophoretic graph showing a major peak between 250-700bp corresponding to fragmented PCR products in B which was pooled and sheared by ultra-sonication. D, SARS-CoV-2 sequence coverage comparison using target enrichment and shotgun sequencing methods in one sample (P1/UAE/2020). *top,* sequencing coverage across the SARS-CoV-2 genomic positions using shotgun transcriptome sequencing (average coverage ~200x); *bottom*, sequencing coverage across SARS-CoV-2 genome (from same patient sample P1/UAE/2020) using target enrichment (average coverage ~1,400x). E, an example of *De novo* assembly of the viral genome isolated from patient P1/UAE/2020 shows clear overlap with the SARS-CoV-2 reference genome.

To enrich viral sequences and minimize sequencing cost and data storage issues addressed below, we describe an alternative approach where the entire SARS-CoV-2 genome is first amplified using 26 overlapping primer sets each yielding around 1.5kb long inserts (**Figures 2A**, **3B** and **Supplemental Table 1**). All inserts were then pooled and fragmented to 250-750bp inserts which were then prepared for short read next generation sequencing (**Figures 2A** and **3C**).

RNA extracted from eight COVID-19 patients, which were first sequenced by shotgun transcriptome (**Table 1** and **Supplemental Table 2**), were sequenced using the enrichment protocol. As expected, we observed significant enhancement in virus detection using this protocol where, on average, 99.3% of the reads now mapped to the SARS-CoV-2 genome leading to tenfold increase in coverage relative to shotgun transcriptome (avg. 440x versus 45.5x, respectively) despite generating a two hundred fold less sequencing data (avg. 0.02Gb versus 4.28Gb, respectively, **Table 2** and **Figure 3D**).

**Table 2.**
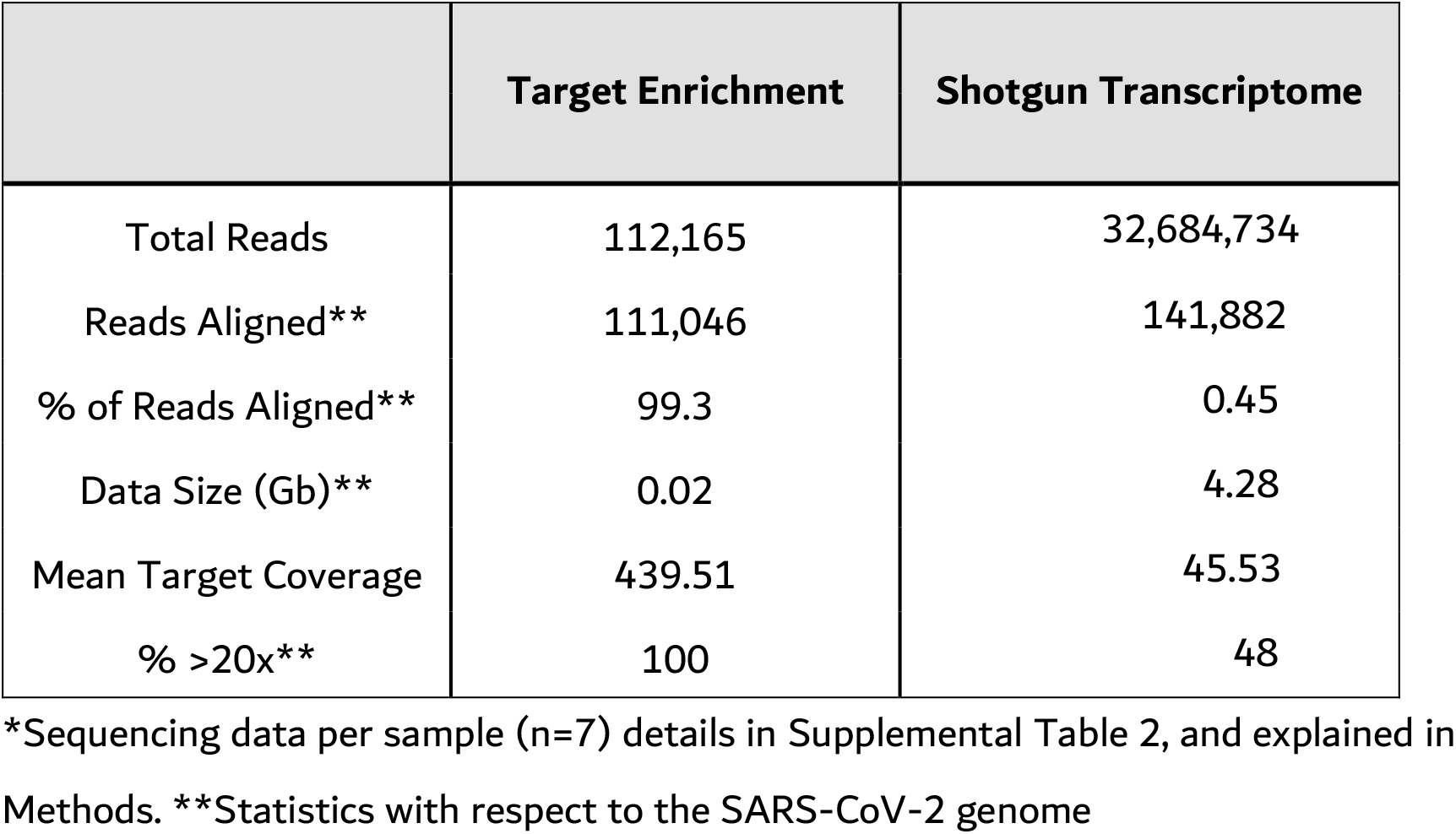
Comparison of sequencing statistics between target enrichment and shotgun transcriptome for COVID-19 patients*

|  | Target Enrichment | Shotgun Transcriptome |
| --- | --- | --- |
| Total Reads | 112,165 | 32,684,734 |
| Reads Aligned** | 111,046 | 141,882 |
| % of Reads Aligned** | 99.3 | 0.45 |
| Data Size (Gb)** | 0.02 | 4.28 |
| Mean Target Coverage | 439.51 | 45.53 |
| % >20x** | 100 | 48 |

\*Sequencing data per sample (n=7) details in Supplemental Table 2, and explained in Methods. \*\*Statistics with respect to the SARS-CoV-2 genome

### Cost, data storage and scalability

On average, 37x coverage per 1Gb of sequencing data was generated using shotgun sequencing (**Table 1**) compared to ~23,000x per 1Gb using target enrichment (**Table 2**) suggesting the latter method is more cost effective and is highly scalable. We calculate the cost of SARS-CoV-2 full genome sequencing to be ~$87 per sample when sequencing 96 samples in a batch at 400x using the target enrichment method. The number of samples in a batch can be doubled (196) while maintaining a low cost (~$104) and a very high coverage of 40,000x per sample (**Table 3**). On the other hand, the cost of sequencing one sample at a much lower coverage (50x) using the shotgun method is $403, while increasing sequencing coverage more than doubled the cost ($1735 at 100x and $1060 at 200x) (**Table 3**). However, using higher throughput sequencing can significantly lower the cost of shotgun sequencing to $232 for 62 samples in a batch at 200x per sample. Nonetheless, using a similar throughput, the per sample cost of enrichment sequencing is $108 for 196 samples in a batch where each sample receives significantly more coverage (~40,000x) (**Table 3**). Therefore, target enrichment sequencing is still more cost-effective and scalable than shotgun transcriptome sequencing even at higher sequencing throughputs.

**Table 3.**
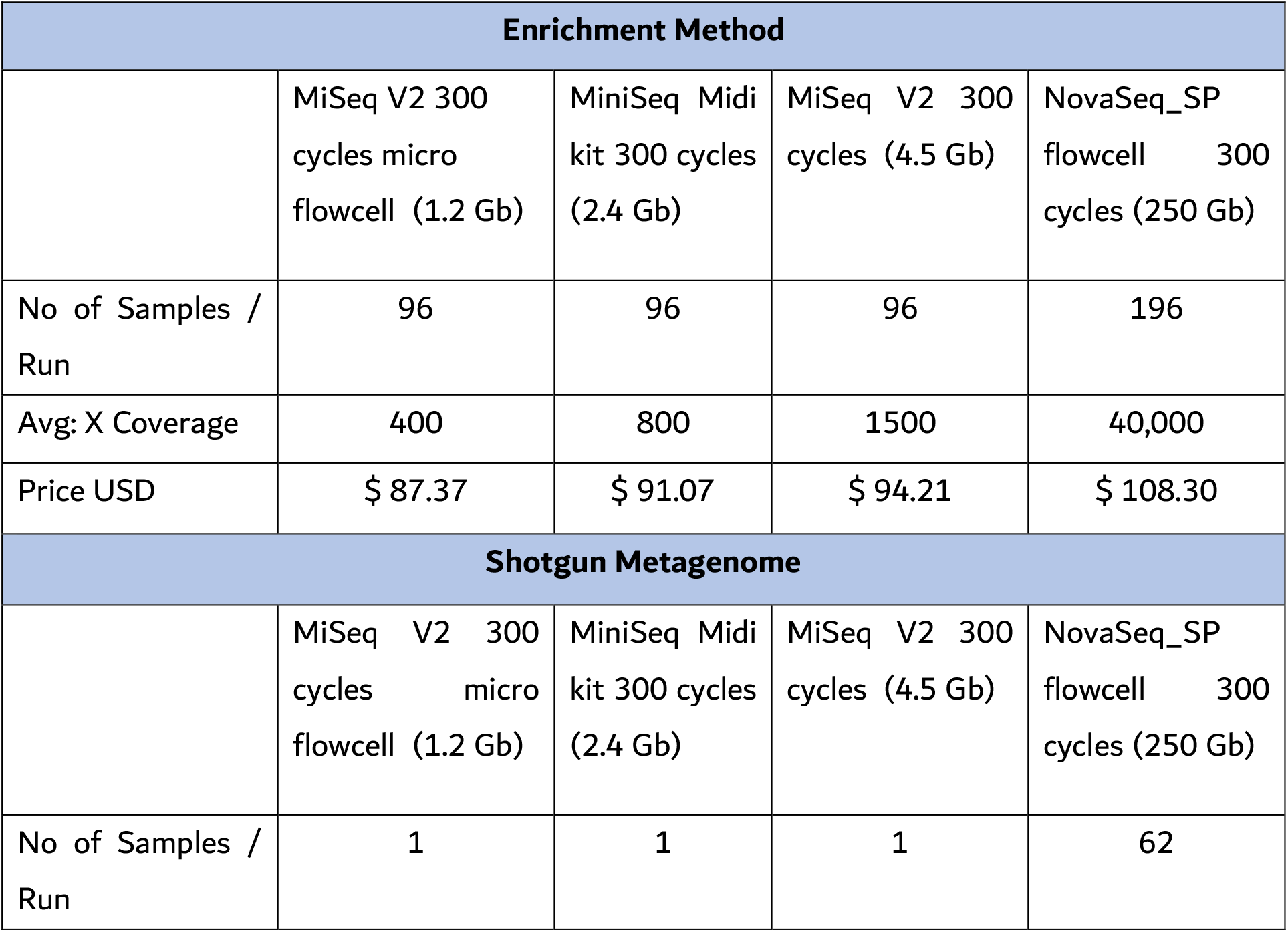
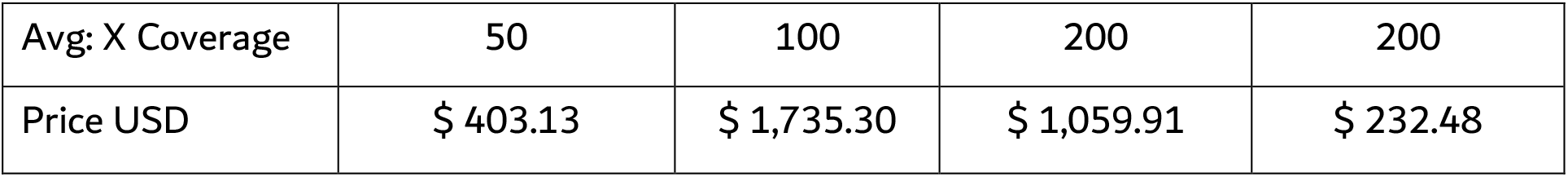
Cost of SARS-CoV-2 whole genome sequencing using Enrichment or shotgun sequencing at different throughputs

| Enrichment Method |  |  |  |  |
| --- | --- | --- | --- | --- |
|  | MiSeq V2 300<br>cycles micro<br>flowcell (1.2 Gb) | MiniSeq Midi<br>kit 300 cycles<br>(2.4 Gb) | MiSeq V2 300<br>cycles (4.5 Gb) | NovaSeq_SP<br>flowcell 300<br>cycles (250 Gb) |
| No of Samples /<br>Run | 96 | 96 | 96 | 196 |
| Avg: X Coverage | 400 | 800 | 1500 | 40,000 |
| Price USD | \$ 87.37 | \$ 91.07 | \$ 94.21 | \$ 108.30 |
| Shotgun Metagenome |  |  |  |  |
|  | MiSeq V2 300<br>cycles micro<br>flowcell (1.2 Gb) | MiniSeq Midi<br>kit 300 cycles<br>(2.4 Gb) | MiSeq V2 300<br>cycles (4.5 Gb) | NovaSeq_SP<br>flowcell 300<br>cycles (250 Gb) |
| No of Samples /<br>Run | 1 | 1 | 1 | 62 |
| Avg: X Coverage | 50 | 100 | 200 | 200 |
| Price USD | \$ 403.13 | \$ 1,735.30 | \$ 1,059.91 | \$ 232.48 |

Another factor impeding scalability of the shotgun approach is data storage. Even with higher throughput sequencing (NovaSeq SP flowcell), shotgun sequencing requires an allocation of 1TB of data for ~250 sequenced samples. On the other hand, with 1TB of data, a total of around 80,000 samples can be sequenced using the enrichment method and the MiSeq Micro flowcell (**Table 3**). Therefore, long term data storage allocations, and cost, are significantly higher, and perhaps formidable, when using the shotgun sequencing approach.

### Genomic surveillance of SARS-CoV-2 origin

To illustrate the utility of SARS-CoV-2 whole genome sequencing, we tracked the origin(s) of the virus in seven patients (UAE/P1/2020, L0287, L1189, L4711, L5857, L6841, L9119) by comparing their assembled sequences to virus strains (n=25) identified during the early phase of the pandemic, between January 29 and March 18 2020, in the UAE (4). All seven patient samples were collected between March 28 and April 5 2020, and are therefore good candidates to determine whether transmissions were community-based due to the previously documented 25 strains or were independent external introductions.

Multiple sequence alignment and phylogenetic analysis (**Figures 2B** and **4**) showed that the new isolates from patients L0287, L4711, L1189, and L5857 clustered with earlier strains of Iranian origin (or clade A3), while that from patient L9119 belonged to the early European (or clade A2a) cluster (**Figure 4**). This information suggests that transmissions for all those five patients were most likely community-based, which we then confirmed from patient medical records where no recent travel history was reported by any of those individuals.

**Figure 4.**
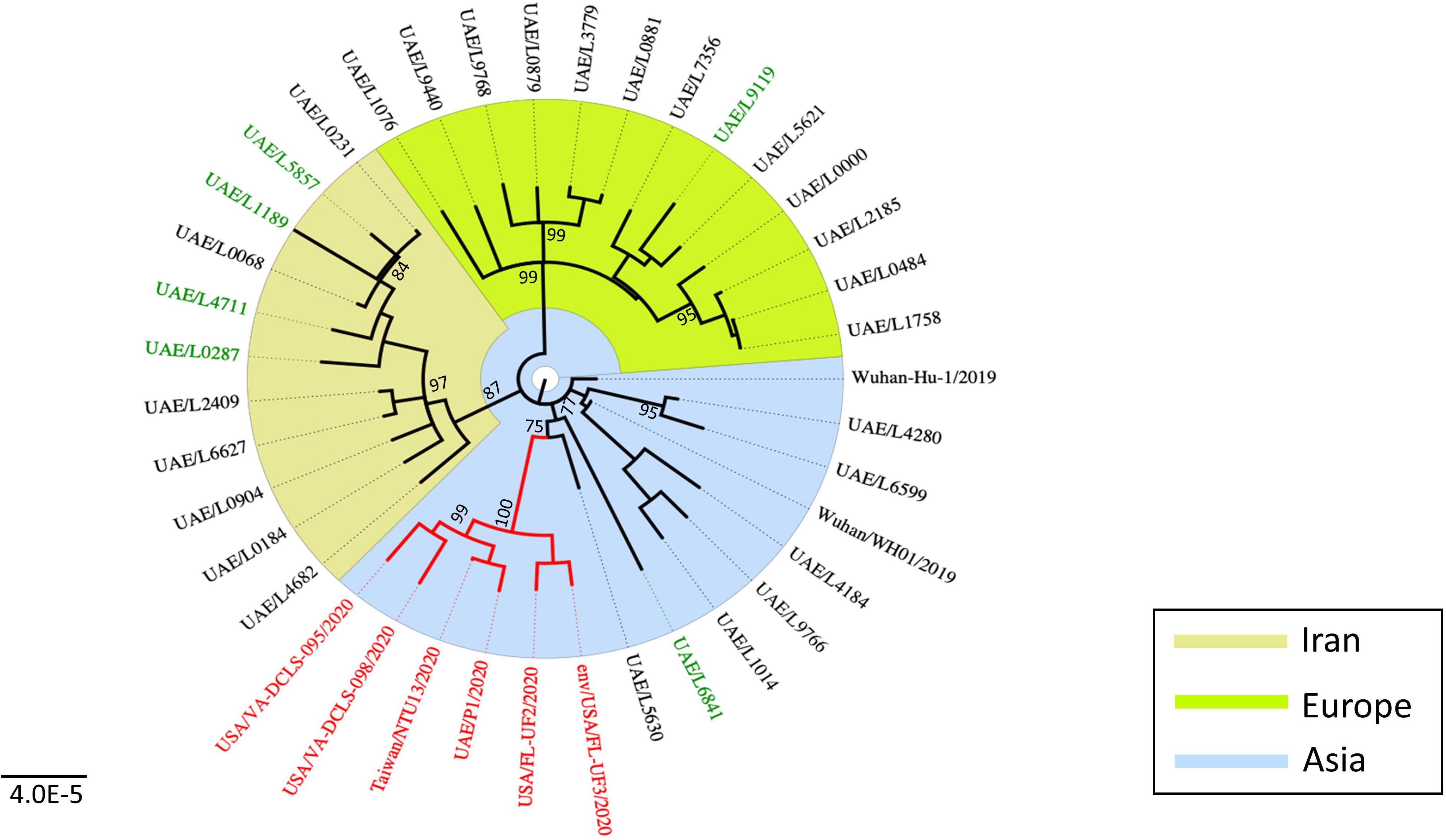
Phylogenetic relationships of SARS-CoV-2 isolates from new patients in this study and previous ‘early’ patients in the UAE, and other countries. A maximum likelihood phylogeny of 37 SARS-CoV-2 genomes (7 obtained in this study (Supplemental Table 2), 5 downloaded from GISAID database (https://www.epicov.org/), and 25 genomes from early patients in UAE (4)). Bootstrap values >70% supporting major branches are shown. The previous European, Iranian, and Asian clusters are highlighted. The 5 non-UAE isolates were selected based on a BLAST search against GISAID database (last accessed 11 May 2020) and high similarity to the P1/UAE/2020 isolate (all red branches). Scale bar represents number of nucleotide substitutions per site. UAE = United Arab Emirates. GISAID = Global Initiative on Sharing All Influenza Data.

SARS-CoV-2 isolates from patients P1/UAE/2020 and L6841 were, on the other hand, closer to the earliest Asian strains, which are more diverse due to fewer but distinct mutations (**Figure 4**). Hence, with the available sequencing data it is challenging to ascertain whether the P1/UAE/2020 and L6841 transmissions were community-based or due to early independent introductions. However, patient L6841 did not have any recent travel history before symptoms onset, suggesting the case for community-based transmission related to an Asian strain. On the other hand, travel history in patient P1/UAE/2020 was not known and the corresponding isolate appeared to match closely to five other strains from the United States and Taiwan (**Figure 4**). Therefore, transmission in patient P1/UAE/2020 was unlikely to be community-based from the early 25 strains (4), but rather due to an independent travel-related introduction of the virus.

## Discussion

Genomics-based SARS-CoV-2 population-based surveillance is a powerful tool for controlling viral transmission during the next phase of the pandemic. Therefore, it is important to devise efficient methods for SARS-CoV-2 genome sequencing for downstream phylogenetic analysis and virus origin tracking. Towards this goal, we describe a cost-effective, robust, and highly scalable target enrichment sequencing approach, and provide an example to demonstrate its utility in characterizing transmission origin.

Our target enrichment protocol is amplicon-based for which oligonucleotide primers can be easily ordered by any molecular laboratory. Next generation sequencing (NGS) has also become largely accessible to most labs, and in our protocol we show that highly affordable, low throughput sequencers, such as the Illumina MiSeq system, can be used efficiently to sequence up to 96 samples at 400x coverage each at a cost of $87 per sample (**Table 3**). This cost is likely comparable to RT-PCR testing for the virus.

One possible limitation is the use of ultra-sonication for fragmentation of PCR products after SARS-CoV-2 whole genome amplification. Several labs might lack sonication systems due to accessibility and affordability issues. In such situations, our protocol can be easily modified to use enzymatic fragmentation instead provided by commercial kits, such as the Agilent SureSelect^QXT^ kit. Furthermore, we have added M13 tails to all our primer sets making them amenable to Sanger sequencing for those labs not equipped with NGS. However, with this approach, manual analysis of sequencing data limits scalability of the approach.

Upon sequence generation, the bioinformatics analysis can be performed using open source scripts. Labs without bioinformatics expertise or support can use online tools (INSaFlu: https://insaflu.insa.pt/; Genome Detective: https://www.genomedetective.com/app/typingtool/virus/) (7,8) which can take raw sequencing (Fastq) files to assemble viral genomes, and to perform multiple sequence alignment and phylogenetic analysis for virus origin tracking. In addition, the described approach does not require significant data storage or computational investment as shown by our cost, data, and scalability calculations (**Table 3**).

Our phylogenetic analysis demonstrates how SARS-CoV-2 genomic sequencing can be used to track origins of virus transmission. However, data should be carefully interpreted, and should be combined with other epidemiological information (such as travel history) to avoid inaccurate conclusions. The major limitation facing genomic-based SARS-CoV-2 surveillance includes the lack of virus sequencing data representing most strains in any country. Nonetheless, SARS-CoV-2 strains are continuously being sequenced by government, private, and academic entities all over the world, and the sequencing data is being shared publicly. This proliferation of sequencing datasets will empower genomic-based surveillance of the virus, and the availability of cost effective sequencing options, like the one described in this study, will contribute to democratizing SARS-CoV-2 sequencing and data sharing.

In summary, we show that SARS-CoV-2 whole genome sequencing is a highly feasible and effective tool for tracking virus transmission. Genomic data can be used to determine community-based versus imported transmissions, which can then inform the most appropriate public health decisions to control the pandemic.

## Acknowledgements

Authors would like to thank members of the Microbiology Laboratory, Latifa Women and Children Hospital, Dubai Health Authority and Al Jalila Children’s Specialty Hospital Genomics Center for supporting SARS-CoV-2 diagnostic testing and for arranging samples used in this study.

