## Appendix I for "SARS-CoV-2 Whole Genome Amplification and Sequencing for Effective Population-Based Surveillance and Control of Viral Transmission"

**SARS-CoV-2 Genome Sequencing using Enrichment method**

| **Reagents, Instruments & Consumables** | |
| --- | --- |
| 1 | Qubit RNA XR Assay Kit - Thermo Fisher Scientific |
| 2 | QuantiTect Reverse Transcription Kit - QIAGEN |
| 3 | Platinum™ SuperFi™ PCR Master Mix |
| 4 | Agencourt AMPure XP beads, 60 mL A63881 |
| 5 | SureSelect XT enrichment kit for illumina |
| 6 | Nuclease Free Water |
| 7 | SARS-CoV-2 M13 tagged primers |
| 8 | DNase/RNase SAP |
| 9 | Micropipette and Tips (10,200,1000ul) |
| 10 | PCR grade 96 well 0.3ml plates |
| 11 | Adhesive Plate seals |
| 12 | Cold blocks |
| 13 | Thermal Cycler |
| 14 | Ethanol |
| 15 | NaOH |
| 16 | Lonza 2.2% Gel |

**Workflow**

Viral RNA Extraction

gDNA Removal and cDNA Synthesis

cDNA Amplification using Genome specific m13 tagged Primers

Post PCR Cleanup using Ampure XP beads

Quantify and Pool the samples

SureSelect XT AGC Custom Protocol

Pool the libraries and Load to Sequencer

Data Analysis

1. **Viral RNA extraction**

Viral RNA extraction performed using QIAGEN QIAmp Viral Mini Kit PN 52904. And QC assessed using Qubit RNA XR Assay Kit - Thermo Fisher Scientific.

2. **gDNA Removal and cDNA Synthesis**

gDNA removal and cDNA synthesis is performed as per the QuantiTect Reverse Transcription Kit – QIAGEN kit protocol.

RNA Requirement – 10-1000 ng

2.1. Thaw template RNA on ice. Thaw gDNA Wipeout Buffer, Quantiscript Reverse Transcriptase, Quantiscript RT Buffer, RT Primer Mix, and RNase-free water at room temperature (15–25°C).

Mix each solution by flicking the tubes. Centrifuge briefly to collect residual liquid

from the sides of the tubes, and then store on ice.

2.2. Prepare the genomic DNA elimination reaction (**2 reactions for one sample**) on ice according to Table 1.

Mix and then store on ice.

**Table 1. Genomic DNA elimination reaction components**

| **Component** | **Volume/reaction** | **Final concentration** |
| --- | --- | --- |
| gDNA Wipe-out Buffer, 7x | 2 µl | 1x |
| Template RNA Variable | Variable (up to 1 µg*) |  |
| RNase-free water | Variable |  |
| **Total volume** | 14 µ**l** |  |

2.3. Incubate for 2 min at 42°C. Then place immediately on ice.

*Note: Do not incubate at 42°C for longer than 10 min.*

2.4. Prepare the reverse-transcription master mix on ice according to **Table 2**.

Mix and then store on ice. The reverse-transcription master mix contains all

components required for first-strand cDNA synthesis except template RNA.

QuantiTect Reverse Transcription Handbook 03/2009 13

**Table 2. Reverse-transcription reaction components**

| **Component** | **Volume/reaction** | **Final concentration** |
| --- | --- | --- |
| **Reverse-transcription master mix** | | |
| Quantiscript Reverse Transcriptase | 1 µl |  |
| Quantiscript RT Buffer, 5x | 4 µl | 1x |
| RT Primer Mix | 1 µl |  |
| **Template RNA** | | |
| Entire genomic DNA elimination reaction (step 3) | 14 µl (add at step 5) |  |
| **Total volume** | 20 µ**l** |  |

2.5. Add template RNA from step 3 (14 µl) to each tube containing reverse transcription master mix. Mix and then store on ice.
2.6. Incubate for 15 min at 42°C.
2.7. Incubate for 3 min at 95°C to inactivate Quantiscript Reverse Transcriptase.

3. **Amplification using, Genome specific M13 tagged primers.**
   3.1. Make a PCR master mix as mentioned in **Table 3**.

| **Component** | **Volume** |
| --- | --- |
| Platinum™ SuperFi™ PCR Master Mix | 12.5ul x 27 Rxns |
| GC Enhancer | 5ul x 27 Rxns |
| Nuclease free water | 1.75ul x 27 Rxns |
| Final cDNA from RT | 1.48 |
| **Total** | **20.73 ul** |

3.2. Aliquot 20ul of the master mix to 26 wells of a plate
3.3. Add 1.25ul of 10pM Forward and reverse primers to separate wells of the reaction plate containing master mix.
3.4. Seal the plate, Pulse Vortex and centrifuge 290g for 1min
3.5. Place in a Thermal Cycler and run the following Protocol.

| **Stage** | | **Temperature (°c)** | **Time** | **Cycles** |
| --- | --- | --- | --- | --- |
| 1 | Initial Denaturation | 98 | 60 Sec | 1 |
| 2 | Denaturation | 98 | 17 Sec | 27 |
| 3 | Annealing | 57 | 20 Sec |  |
| 4 | Extension | 72 | 1min.5 Sec |  |
| 5 | Final Extension | 72 | 10 min | 1 |
| 6 | Hold | 4 | ∞ | ∞ |

3.6. Once the PCR is done, Run 2ul of the product in an Agarose Gel.

Gel QC

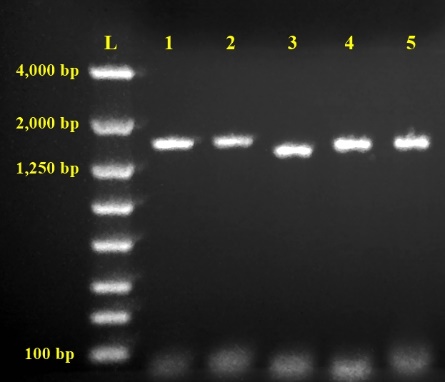

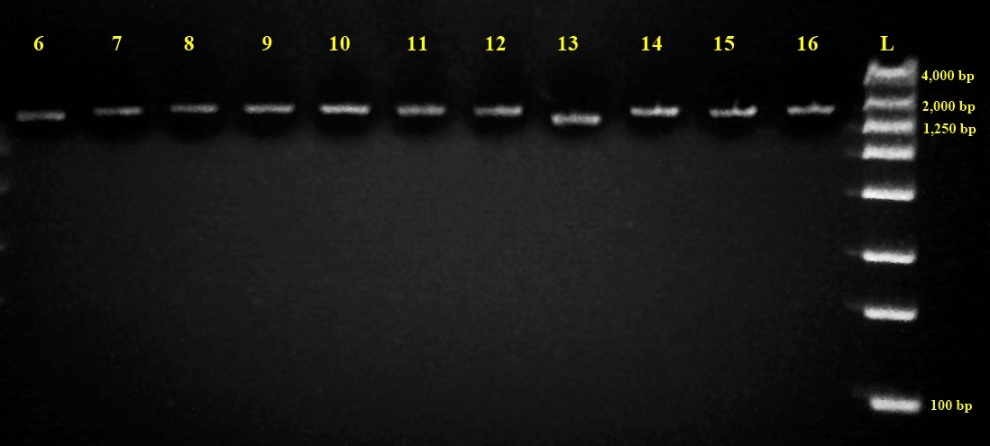

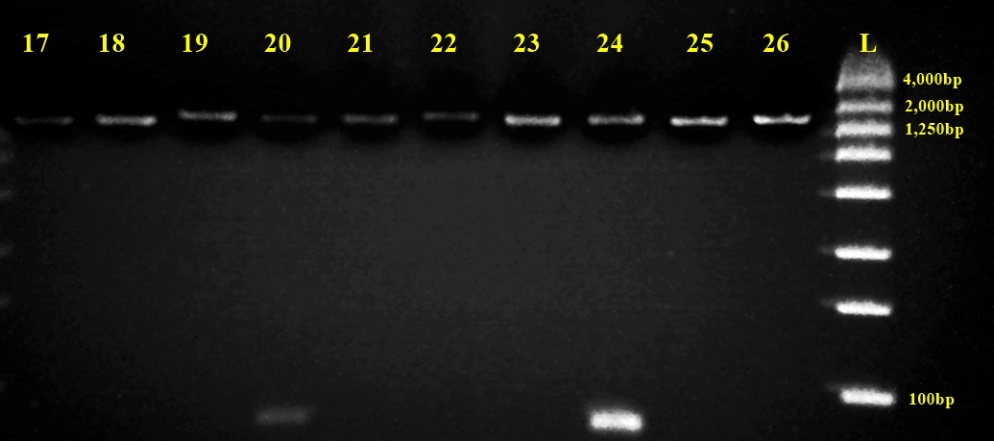

4. **Purify Amplified products with Ampure XP beads**
   4.1. Shake the Agencourt AMPure XP bottle to re-suspend any magnetic particles that may have settled. The volume of Agencourt AMPure XP for a given reaction can be derived from the following equation:

Volume of Agencourt AMPure XP per reaction = 1.8 x Reaction Volume.

Add the Agencourt AMpure XP according to the sample reaction volume presented in **Table 4**.

| **Sample Reaction Volume (µl)** | **AMPure XP Volume (µl)** |
| --- | --- |
| 10 | 18 |
| 20 | 36 |
| 50 | 90 |
| 100 | 180 |

Table 1: AMPure XP to Sample Reaction volume Chart.

4.2. Mix reagent and sample thoroughly by pipetting 10 times. Let the mixed samples incubate for 5 minutes at room temperature for maximum recovery.

Note: This step binds DNA fragments 100bp and larger to the magnetic beads. Mixing by pipetting is preferable to vortexing as it tends to be more reproducible. The color of the mixture should appear homogenous after mixing.

4.3. Place the reaction plate onto a 96 well plate Magnet stand for 2 minutes to separate beads from the solution. Note: Wait for the solution to clear before proceeding to the next step.
4.4. While the reaction plate is situated on the 96 well plate Magnet stand, aspirate the cleared solution from the reaction plate and discard. Leave 5µl of supernatant behind, to prevent the beads from being drawn out with the supernatant. Do not disturb the ring of separated magnetic beads.
4.5. Dispense 200µl of 70% Ethanol to each well of the reaction plate that is situated on the 96 well magnetic plate. Incubate for 30 seconds at room temperature. Aspirate out all the ethanol and discard.

**Repeat a total of two washes**. A dry time of 7 minutes is favorable to ensure all traces of ethanol are removed. Do not over dry the beads (bead ring appears cracked if over dried), as this will significantly decrease the elution efficiency.

*Note: The beads are not drawn out easily when in alcohol, so it is not necessary to leave any supernatant behind.*

4.6. Remove the reaction plate from the magnet plate, and then add 10µl of nuclease free water to each well of the reaction plate and mix pipette 10 times. Incubate for 2 minutes.

4.7. Place the reaction plate onto a 96 well plate Magnet stand for 2 minutes to separate beads from the solution.

4.8. Transfer the eluate to a new plate, or tubes, in case of few samples and discard the beads.

5. **Quantify and Pool the samples**
   5.1. Quantify the purified PCR product using a nanodrop**.** Dilute the samples to the same conc and Pool them into one tube.
6. **SureSelect XT AGC Customized protocol**

The Sureselect XT assay optimized to work with low input initial DNA than the recommended 3μg, and the library can be run in Illumina NovaSeq 6000 and MiSeq sequencer using paired-end sequencing with a read length of 2x151 base pairs. The protocol has been tested with total DNA concentration ranging from 200-800ng.

This protocol was revised from the Agilent SureSelect XT user guide.

### Overview of the Workflow

Final Libraries

Purified PCR product (200-800ng)

Shear DNA using Covaris

DNA Fragment size (250-700bp)

Adaptor-Tagged DNA libraries

Indexing libraries

indexIndexing libraries tagged final libraries

Sequencing

### Required Consumables and Equipment

| **Consumables** | **Equipment** |
| --- | --- |
| Ampure XP Beads | Thermo Veriti Thermal cycler |
| Dynabeads MyOne Streptavidin T1 magnetic beads P/n | Covaris LE220 plus |
| 100% Ethanol, molecular biology grade | Vortex Mixer |
| Herculase II Fusion DNA Polymerase (includes dNTPs and 5× Buffer) P/n 600677 | Plate centrifuge |
| Qubit dsDNA HS Assay Kit P/n Q32851 | Mini centrifuge for 1.5 and 2ml tubes |
| Qubit dsDNA BR Assay Kit P/n Q32853 | illumina NovaSeq 6000 |
| Nuclease free water (Not DEPC treated) | Nanodrop |
| SureSelectXT Clinical Research Exome V2 | Qubit Fluorometer |
| Agilent TapeStation 4200 - High Sensitivity D1000 ScreenTape and D1000 ScreenTape | TapeStation 4200 |
| Agilent TapeStation 4200 -High Sensitivity D1000 and D1000 Reagents | 0.2, 10, 20, 100, 200 and 1000μl Single channel pipettes and Multi-channel pipettes |
| 0.3ml 96 well PCR plate - BioRad | Easypet 3 |
| 0.8ml 96 well MIDI plate - Thermo | 96 well Magnetic stand |
| 0.2ml 8 well strips and cap | 96 well Plate shaker |
| 1.5ml, 2ml Micro centrifuge tubes |  |
| 15ml and 50ml Conical centrifuge tubes |  |
| 10, 20, 100, 200, 1000μl, barrier Pipette tips |  |
| Adhesive clear sealing film for PCR plates |  |
| SureSelect XT Library Prep Kit ILM (96 Samples) P/n 5500-0133 |  |
| SureSelect Target Enrichment-Box 1 P/n 5190-8646 |  |
| SureSelect Target Enrichment Kit ILM Indexing Hyb Module Box 2 P/n 5190-4456 |  |
| NovaSeq Reagent kit 300 cycles (SP, S1 & S4) |  |
| Covaris - 8 microTUBE Strip (130 µL) (12) (PN 520053) |  |
| 0.5ml Qubit assay tubes |  |
| Reagent reservoirs |  |
| Tapestation 4200 loading tips |  |

#### A. **DNA Requirements**

A minimum of 200-800ng in a total volume of 55µl is required to start the assay.

#### 6.1. **Dilution and Shearing of genomic DNA**

6.1.1 Dilute 200-800ng of high-quality DNA with nuclease free water in a 96-well plate or 8-strip tube to a total volume of **55 μL**.
6.1.1.
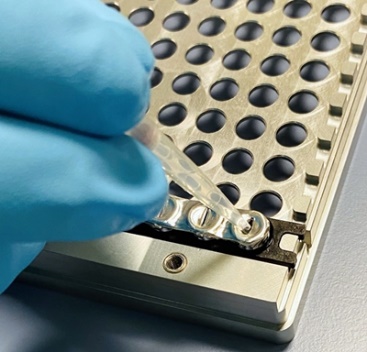
 Pierce the foil seal on the Covaris glass microTube with 200μL tip.
6.1.2. Use a tapered pipette tip to slowly transfer the **55 μL** DNA sample through the foil seal (Avoid bubbles into the bottom of the tube) to the microTUBE. (Pulse Centrifuge if bubbles are present).

**** A second technologist must verify the samples and their well position**

6.1.3. Once all the tubes are filled with sample, use a new foil seal to seal the strip tubes properly.
6.1.4. Secure the microTUBE strip in the plate holder and screw on the lid of Covaris plate tightly.
6.1.5. Shear the DNA with the following settings (AJCH-Covid-Amplicon shear) in a Covaris instrument to generate a target fragment size of 250-700bp.

**Table 1:** AJCH-Covid-Amplicon shear

| **Setting** | **Value** |
| --- | --- |
| Duty Factor | 20% |
| Peak Power | 150 Watts |
| Cycles per Burst | 900 |
| Treatment Time | 320 seconds |
| Avg Power | 30 Watts |
| Bath Temperature | 20° C |

6.1.6. After shearing, while keeping the strip tube on, insert a pipette tip through the foil seal, then slowly remove the sheared DNA (careful not to Overflow).
6.1.7. Transfer each sheared DNA sample (approximately 55 μL) to a separate well of a 96-well MIDI plate.

#### 6.2. **Purify the sample using AMPure XP beads**

6.2.1. Let the AMPure XP beads come to room temperature for at least **30 minutes**. Do not freeze the beads at any time.
6.2.2. Prepare **400μL of 70% ethanol** per sample, for use in step 2.8.
6.2.3. Vortex the bead suspension well so that the reagent appears homogeneous and consistent in color.
6.2.4. **Add 90μL of homogeneous AMPure XP beads** to each sheared DNA sample (approximately 55μL) in the MIDI plate. Pipette up and down 10 times to mix.
6.2.5. Incubate samples for **5 minutes** at **room temperature**.
6.2.6. Put the plate into a magnetic separation device. Wait for the solution to clear (approximately 3 to 5 minutes).
6.2.7. Keep the plate in the magnetic stand. Carefully remove and discard the cleared solution (supernatant) from each well. Do not touch the beads pellet while removing the solution.
6.2.8. Continue to keep the plate in the magnetic stand while dispensing **200μL of 70% ethanol** in each sample well. Use fresh 70% ethanol for optimal results.
6.2.9. Wait for 1 minute to allow any disturbed beads to settle, then remove the ethanol.
6.2.10. Repeat step 8 to step 9 once (2X Ethanol wash).
6.2.11. **After ethanol removal, air dry** the samples for **7 minutes at room temperature** while still on magnet stand.
6.2.12. Take the sample tubes(s) **OFF** the magnet and Add **30μL nuclease-free water** to each sample well.
6.2.13. Pipette up and down 10 times to mix (Ensure the mixture is homogenous).
6.2.14. Incubate for **2 minutes** at room temperature.
6.2.15. Put the plate **ON** the magnetic stand and leave for 2 to 3 minutes, until the solution is clear.
6.2.16. Remove the cleared supernatant (**approximately 28μL**) to a fresh PCR plate well. Discard the beads at this time.

**Stopping Point If not continuing to the next step, seal the plate and store at –4°C. If Continuing NEXT DAY store at 4°C**

#### 6.3. **Assess the quality of the fragmented product using Agilent High Sensitivity D1000 ScreenTape Assay**

6.3.1. Allow High Sensitivity D1000 Reagents (5067-5585) to equilibrate at room temperature for 30 minutes.
6.3.2. Launch the Agilent 4200 TapeStation Controller Software.
6.3.3. Flick the High Sensitivity D1000 ScreenTape device (5067-5584) and load it into the 4200 TapeStation instrument.
6.3.4. Place loading tips (5067-5598) into the Agilent 4200 TapeStation instrument.
6.3.5. Vortex reagents and spin down before use.
6.3.6. Prepare ladder:
- For 1 – 15 samples: pipette 2μL High Sensitivity D1000 Sample Buffer and 2μL High Sensitivity D1000 Ladder at position A1 in a tube strip (401428).
- For 16 or more samples: pipette 15μL High Sensitivity D1000 Sample Buffer and 15μL High Sensitivity D1000 Ladder at position A1 in a tube strip.
6.3.7. For each sample, pipette 2μL High Sensitivity D1000 Sample Buffer and 2μL DNA sample in a well plate (5042-8502) or a tube strip (401428).
6.3.8. Apply foil seal (5067-5154) to sample well plate and caps (401425) to tube strips with ladder or sample.
6.3.9. Mix liquids in sample and ladder vials using the IKA vortex at 2000 rpm for 1 min.
6.3.10. Spin down to position the sample and ladder at the bottom of the well plate and tube strip.

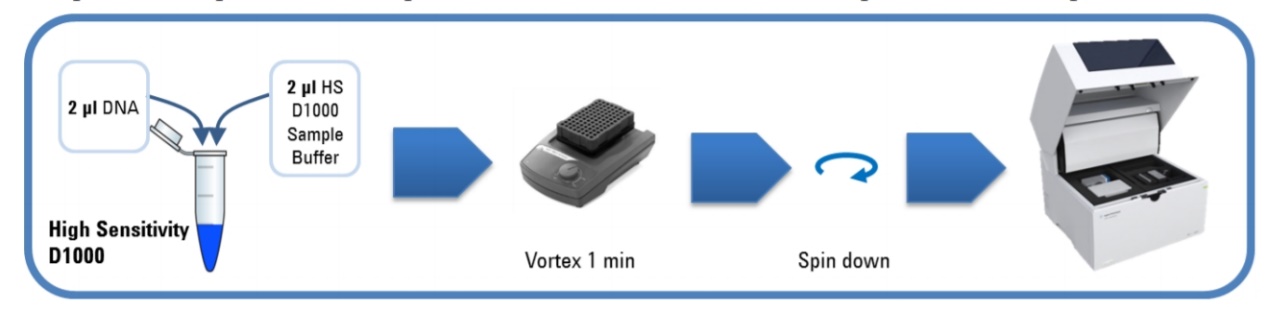

6.3.11. Load samples into the Agilent 4200 TapeStation instrument. Carefully remove caps of tube strips.
6.3.12. Place ladder in position A1 on tube strip holder in the 4200 TapeStation instrument.
6.3.13. Select required sample positions on the 4200 TapeStation Controller Software.
6.3.14. Click Start.
6.3.15. The Agilent TapeStation Analysis Software opens after the run and displays results.
6.3.16. The estimated Avg fragment size of the product is from 250-500bp.

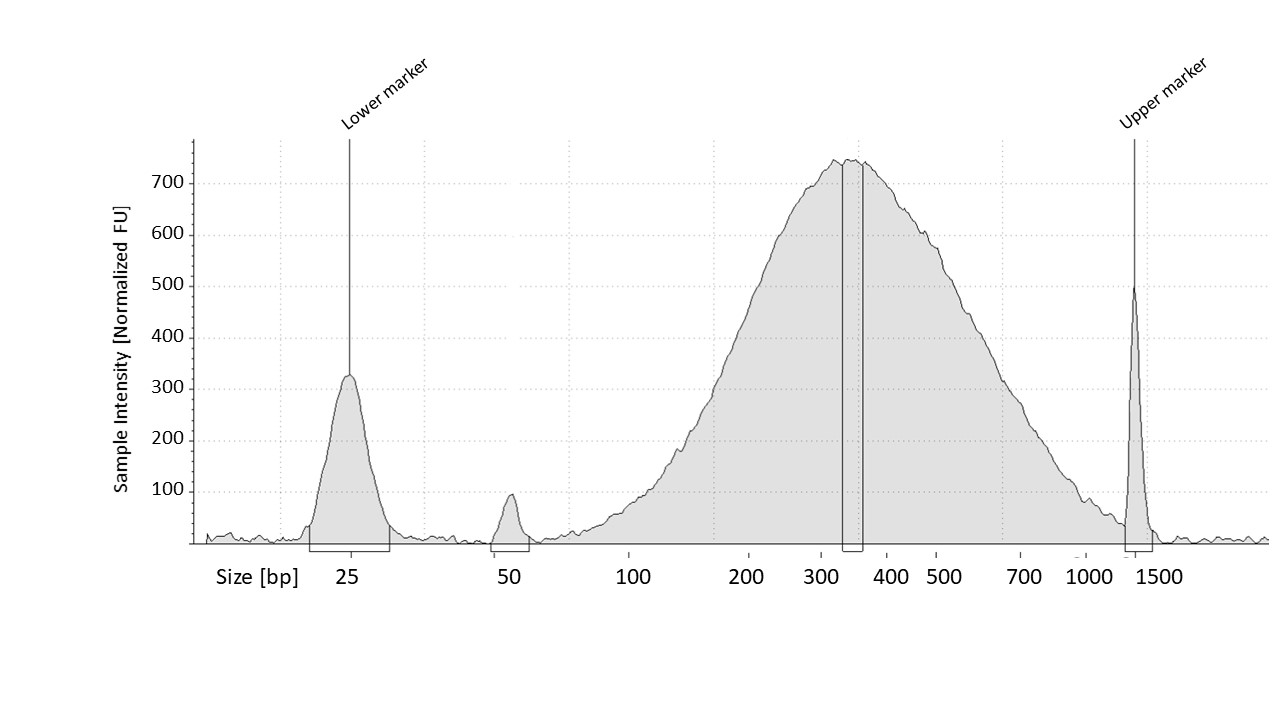

Figure Showing TapeStation profile of fragmented DNA with an average size of 360bp

#### 6.4. **End Repair**

6.4.1. Prepare the appropriate volume of End Repair master mix on ice, as described in **Table 2**. Mix well on a vortex mixer.

**Table 2:** End repair master mix preparation.

| **Reagent** | **Volume for 1 reaction** | **Volume for 16 reactions (includes excess)** |
| --- | --- | --- |
| Nuclease-free water | 35.2μL | 580.8μL |
| 10× End Repair Buffer (clear cap) | 10μL | 165μL |
| dNTP Mix (green cap) | 1.6μL | 26.4μL |
| T4 DNA Polymerase (purple cap) | 1μL | 1μL |
| Klenow DNA Polymerase (yellow cap) | 2μL | 33μL |
| T4 Polynucleotide Kinase (orange cap) | 2.2μL | 36.3μL |
| **Total** | **52μL** | **858μL** |

6.4.2. Vortex Master mix and spin to collect.
6.4.3. Add **52μL** of the master mix to each PCR plate sample well containing purified, sheared DNA. Mix by pipetting up and down.
6.4.4. Incubate the plate in the thermal cycler and run the program in **Table 3**. Do not use a heated lid.

**Table 3:** End-Repair Thermal Cycler Program.

| **Step** | **Temperature** | **Time** |
| --- | --- | --- |
| **Step 1** | 20 °C | 30 minutes |
| **Step 2** | 4 °C | Hold |

#### 6.5. **Purify the sample using AMPure XP beads**

6.5.1. Let the AMPure XP beads come to room temperature for at least **30 minutes**.
6.5.2. Mix the bead suspension well so that the reagent appears homogeneous and consistent in color.
6.5.3. Transfer samples from 96-well plate **(approximately 80μL)** to a fresh 96-well MIDI plate.
6.5.4. Add **150μL** of homogeneous AMPure XP beads to each **80μL** end-repaired DNA sample in the MIDI plate. Pipette up and down 10 times to mix.
6.5.5. Incubate samples for 5 minutes at room temperature.
6.5.6. Put the plate into a magnetic separation device. Wait for the solution to clear (approximately 3 to 5 minutes).
6.5.7. Keep the plate in the magnetic stand. Carefully remove and discard the cleared solution/supernatant from each well. Do not touch the beads/pellet while removing the solution.
6.5.8. Continue to keep the plate in the magnetic stand while dispensing **200μL of freshly-prepared 70% ethanol** in each sample well.
6.5.9. Wait for 1 minute to allow any disturbed beads to settle, then remove the ethanol/supernatant.
6.5.10. Repeat step 8 to step 9 once (2X Ethanol wash)
6.5.11. **After ethanol removal, air dry** the samples for **7 minutes at room temperature** while still on magnet stand.
6.5.12. Take the sample tubes **OFF** the magnet and Add **25μL nuclease-free water** to each sample well.
6.5.13. Pipette up and down times to mix (Ensure the mixture is homogenous).
6.5.14. Incubate for **2 minutes** at room temperature.
6.5.15. Put the plate **ON** the magnetic stand and leave for 2 to 3 minutes, until the solution is clear.
6.5.16. Remove the cleared supernatant (**approximately 24μL**) to a fresh PCR plate well. Discard the beads at this time.

**Optional stopping Point If not continuing to the next step, seal the plate and store at –20°C.**

#### 6.6. **Adenylate the 3’ end of the DNA fragments**

6.6.1. Prepare the appropriate volume of Adenylation master mix on ice, as described in **Table 4**. Mix well on a vortex mixer.

**Table 4:**	Adenylataion of the 3' end of the DNA fragments

| **Reagent** | **Volume for 1 reaction** | **Volume for 16 reactions (includes excess)** |
| --- | --- | --- |
| Nuclease-free water | 11μL | 181.5μL |
| 10× Klenow Polymerase Buffer (blue cap) | 5μL | 82.5μL |
| dATP (green cap) | 1μL | 16.5μL |
| Exo(–) Klenow (red cap) | 3μL | 49.5μL |
| **Total** | **20μL** | **330μL** |

6.6.2. Vortex Master mix and spin to collect.
6.6.3. Add **20μL** of the Adenylation master mix to each end-repaired, purified DNA sample (approximately 24μL).
6.6.4. Mix well by pipetting up and down.
6.6.5. Incubate the plate in the thermal cycler and run the program in **Table 5**. Do not use a heated lid (Leave the lid open).

**Table 5:** dA-Tailing Thermal Cycler Program (50μL total volume).

| **Step** | **Temperature** | **Time** |
| --- | --- | --- |
| **Step 1** | 37 °C | 30 minutes |
| **Step 2** | 4 °C | Hold |

#### 6.7. Purify the sample using AMPure XP beads

6.7.1. Let the AMPure XP beads come to room temperature for at least **30 minutes**. Do not freeze the beads at any time.
6.7.2. Mix the bead suspension well so that the reagent appears homogeneous and consistent in color.
6.7.3. Transfer samples from 96-well plate **(approximately 44μL)** to a fresh 96-well MIDI plate.
6.7.4. Add **80μL** of homogeneous AMPure XP beads to each **44μL** dA-tailed DNA sample in the MIDI plate. Pipette up and down 10 times to mix.
6.7.5. Incubate samples for 5 minutes at room temperature.
6.7.6. Put the plate into a magnetic separation device. Wait for the solution to clear (approximately 3 to 5 minutes).
6.7.7. Keep the plate in the magnetic stand. Carefully remove and discard the cleared solution/supernatant from each well. Do not touch the beads/pellet while removing the solution.
6.7.8. Continue to keep the plate in the magnetic stand while dispensing 200μL of freshly-prepared 70% ethanol in each sample well.
6.7.9. Wait for 1 minute to allow any disturbed beads to settle, then remove the ethanol.
6.7.10. Repeat step 8 to step 9 step once (2X Ethanol wash).
6.7.11. **After ethanol removal, air dry** the samples for **7 minutes at room temperature** while still on magnet stand.
6.7.12. Take the sample tubes **OFF** the magnet and Add **15μL nuclease-free water** to each sample well.
6.7.13. Pipette up and down times to mix (Ensure the mixture is homogenous).
6.7.14. Incubate for **2 minutes** at room temperature.
6.7.15. Put the plate **ON** the magnetic stand and leave for 2 to 3 minutes, until the solution is clear.
6.7.16. Remove the cleared supernatant (**approximately 13.5μL**) to a fresh PCR plate well. Discard the beads at this time **(Since it is low volume, use 10μL multichannel pipette to transfer 6.5μL samples from MIDI plate to PCR 96-well plate twice).**
6.7.17. **Proceed IMMEDIATELY to the Next Step (Step 7: Ligate the paired-end adaptor).**

#### 6.8. Ligate the paired-end adaptor

6.8.1. Prepare the appropriate volume of Ligation master mix, as described in **Table 6**, on ice. Mix well on a vortex mixer.

**Table 6:** Preparation of Ligation master mix.

| **Reagent** | **Volume for 1 reaction** | **Volume for 16 reactions (includes excess)** |
| --- | --- | --- |
| Nuclease-free water | 15.5μL | 255.75μL |
| 5× T4 DNA Ligase Buffer (green cap) | 10μL | 165μL |
| SureSelect Adaptor Oligo Mix (brown cap) | 10μL | 165μL |
| T4 DNA Ligase (red cap) | 1.5μL | 24.75μL |
| **Total** | **37μL** | 610.5.1 **μL** |

6.8.2. Vortex the Master mix and spin to collect.
6.8.3. Add **37μL** of the Ligation master mix to each dA-tailed, purified DNA sample (**13μL**) in the PCR plate wells.
6.8.4. Mix well by pipetting up and down.
6.8.5. Incubate the plate in the thermal cycler and run the program in **Table 7**. Do not use a heated lid (Leave the lid open).

**Table 7:**	Ligation Thermal Cycler Program (50μL total volume).

| **Step** | **Temperature** | **Time** |
| --- | --- | --- |
| **Step 1** | 20 °C | 15 minutes |
| **Step 2** | 4 °C | Hold |

#### 6.9. **Purify the sample using AMPure XP beads**

6.9.1. Let the AMPure XP beads come to room temperature for at least **30 minutes**. Do not freeze the beads at any time.
6.9.2. Mix the bead suspension well so that the reagent appears homogeneous and consistent in color.
6.9.3. Transfer samples from 96-well plate **(approximately 50μL)** to a fresh 96-well MIDI plate.
6.9.4. Add **90μL** of homogeneous AMPure XP beads to each **50μL** each adaptor-ligated DNA sample in the MIDI plate. Pipette up and down 10 times to mix.
6.9.5. Incubate samples for 5 minutes at room temperature.
6.9.6. Put the plate into a magnetic separation device. Wait for the solution to clear (approximately 3 to 5 minutes).
6.9.7. Keep the plate in the magnetic stand. Carefully remove and discard the cleared solution/supernatant from each well. Do not touch the beads/pellet while removing the solution.
6.9.8. Continue to keep the plate in the magnetic stand while dispensing 200μL of freshly-prepared 70% ethanol in each sample well.
6.9.9. Wait for 1 minute to allow any disturbed beads to settle, then remove the ethanol.
6.9.10. Repeat step 8 to step 9 step once. (2X Ethanol wash).
6.9.11. **After ethanol removal, airdry** the samples for **7 minutes at room temperature** while still on magnet stand).
6.9.12. Take the sample tubes **OFF** the magnet and Add **25μL nuclease-free water** to each sample well.
6.9.13. Pipette up and down times to mix. (Ensure the mixture is homogenous).
6.9.14. Incubate for **2 minutes** at room temperature.
6.9.15. Put the plate **ON** the magnetic stand and leave for 2 to 3 minutes, until the solution is clear.
6.9.16. Remove the cleared supernatant (**approximately 23.5μL**) to a fresh PCR plate well. Discard the beads at this time.

**Stopping Point If not continuing to the next step, seal the plate and store at –20°C. If Continuing NEXT DAY store at 4°C**

#### 6.10. **Amplify the adaptor-ligated library**

6.10.1. Prepare the appropriate volume of pre-capture PCR reaction mix, as described in **Table 8**, on ice. Mix well on a vortex mixer.

**Table 8:** Preparation of SureSelect Pre-Capture PCR Reaction Mix.

| **Reagent** | **Volume for 1 reaction** | **Volume for 16 reactions (includes excess)** |
| --- | --- | --- |
| Nuclease-free water | 21μL | 346.5μL |
| SureSelect Primer (brown cap) | 1.25μL | 20.6μL |
| SureSelect ILM Indexing Pre-Capture PCR  Reverse Primer (clear cap) | 1.25μL | 20.6μL |
| 5× Herculase II Reaction Buffer (clear cap) | 10μL | 165μL |
| 100 mM dNTP Mix (green cap) | 0.5μL | 8.25μL |
| Herculase II Fusion DNA Polymerase (red cap) | 1μL | 16.5μL |
| **Total** | **35μL** | 577.5.1 **μL** |

6.10.2. Transfer **23.5μL** of each purified DNA library to a new PCR 96-Well plate.
6.10.3. Combine **35μL** of the PCR reaction mixture prepared in **Table 8** and 15μL of each purified DNA library sample from step 8.17. Add a single DNA library sample to each well of the plate or strip tube.
6.10.4. Mix by pipetting.
6.10.5. Run the thermal cycling program as in **Table 9**.

**Table 9:**	Pre-Capture PCR Thermal Cycler Program (50μL total volume).

| **Step** | **No of cycles** | **Temperature** | **Time** |
| --- | --- | --- | --- |
| Step 1 | 1 | 98 °C | 2 minutes |
| Step 2 | 7 | 98 °C | 30 seconds |
|  |  | 65 °C | 30 seconds |
|  |  | 72 °C | 1 minutes |
| Step 3 | 1 | 72 °C | 10 minutes |
| Step 4 | 1 | 4 °C | Hold |

#### 6.11. Purify the amplified library with AMPure XP beads

6.11.1. Let the AMPure XP beads come to room temperature for at least **30 minutes**. Do not freeze the beads at any time.
6.11.2. Mix the bead suspension well so that the reagent appears homogeneous and consistent in color.
6.11.3. Transfer samples from 96-well plate **(approximately 58μL)** to a fresh 96-well MIDI plate.
6.11.4. Add **105μL** of homogeneous AMPure XP beads to each **58μL** each adaptor-ligated DNA sample in the MIDI plate. Pipette up and down 10 times to mix.
6.11.5. Incubate samples for 5 minutes at room temperature.
6.11.6. Put the plate into a magnetic separation device. Wait for the solution to clear (approximately 3 to 5 minutes).
6.11.7. Keep the plate in the magnetic stand. Carefully remove and discard the cleared solution/supernatant from each well. Do not touch the beads/pellet while removing the solution.
6.11.8. Continue to keep the plate in the magnetic stand while dispensing 200μL of freshly-prepared 70% ethanol in each sample well.
6.11.9. Wait for 1 minute to allow any disturbed beads to settle, then remove the ethanol.
6.11.10. Repeat step 8 to step 9 step once (2X Ethanol wash).
6.11.11. **After ethanol removal, air dry** the samples for **7 minutes at room temperature** while still on magnet stand.
6.11.12. Take the sample tubes **OFF** the magnet and Add **13.5μL nuclease-free water** to each sample well.
6.11.13. Pipette up and down times to mix (Ensure the mixture is homogenous).
6.11.14. Incubate for **2 minutes** at room temperature.
6.11.15. Put the plate **ON** the magnetic stand and leave for 2 to 3 minutes, until the solution is clear.
6.11.16. Remove the cleared supernatant (**approximately 12.5μL**) to a fresh PCR plate well. Discard the beads at this time. **(Use a single channel pipette to move the sample due to low volume).**

### Assess the Quality and Quantity of index-tagged library

#### 6.12. **Quantify each index-tagged library by Qubit™ 1X dsDNA HS Assay**

6.12.1. Set up the required number of 0.5mL tubes for standards and samples. The Qubit™ 1X dsDNA HS Assay requires 2 standards.

Note: Use only thin-wall, clear, 0.5-mL PCR tubes. Acceptable tubes include Qubit™ assay tubes (Cat. No. Q32856)

6.12.2. Label the tube lids.

Note: Do not label the side of the tube as this could interfere with the sample read.

6.12.3. Label the lid of each standard tube correctly. Calibration of the Qubit™ Fluorometer requires the standards to be inserted into the instrument in the right order.
6.12.4. Add the Qubit™ 1X dsDNA 1X buffer to each tube (199µL) such that the final volume is 200µL
6.12.5. Add 10 µL of each Qubit™ standard to the appropriate tube.
6.12.6. Add 1 µL of each user sample to the appropriate tube.

Note: If you are adding 1–2μL of sample, use a P-2 pipette for best results.

6.12.7. Mix each sample vigorously by vortexing for 3–5 seconds.
6.12.8. Allow all tubes to incubate at room temperature for 2 minutes, then proceed to “Read standards and samples” in the Qubit instrument.
6.12.9. Do the quantification twice for one sample and take the Average for calculating the nM concentration of each library.

#### 6.13. **Assess the quality of library using Agilent High Sensitivity D1000 ScreenTape Assay**

6.13.1. Allow High Sensitivity D1000 Reagents (5067-5585) to equilibrate at room temperature for 30 minutes.
6.13.2. Launch the Agilent 4200 TapeStation Controller Software.
6.13.3. Flick the High Sensitivity D1000 ScreenTape device (5067-5584) and load it into the 4200 TapeStation instrument.
6.13.4. Place loading tips (5067-5598) into the Agilent 4200 TapeStation instrument.
6.13.5. Vortex reagents and spin down before use.
6.13.6. Prepare ladder:
- For 1 – 15 samples: pipette 2μL High Sensitivity D1000 Sample Buffer and 2μL High Sensitivity D1000 Ladder at position A1 in a tube strip (401428).
- For 16 or more samples: pipette 15μL High Sensitivity D1000 Sample Buffer and 15μL High Sensitivity D1000 Ladder at position A1 in a tube strip.
6.13.7. For each sample, pipette 2μL High Sensitivity D1000 Sample Buffer and 2μL DNA sample in a well plate (5042-8502) or a tube strip (401428).
6.13.8. Apply foil seal (5067-5154) to sample well plate and caps (401425) to tube strips with ladder or sample.
6.13.9. Mix liquids in sample and ladder vials using the IKA vortex at 2000 rpm for 1 min.
6.13.10. Spin down to position the sample and ladder at the bottom of the well plate and tube strip.

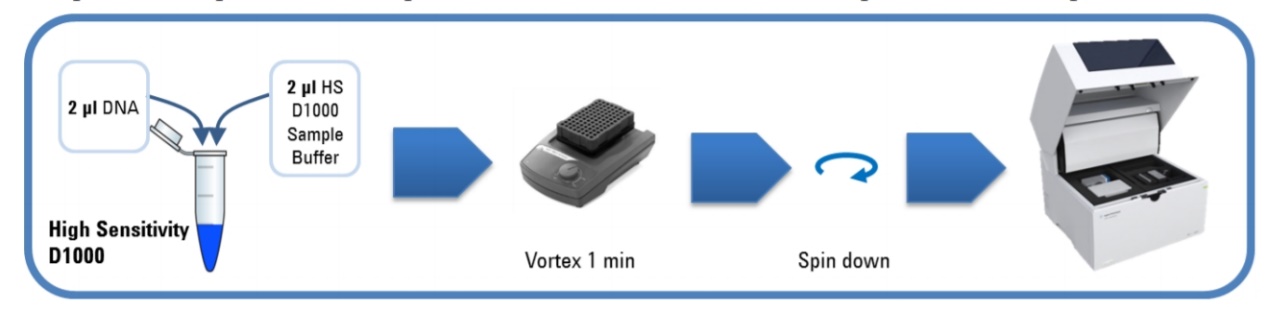

6.13.11. Load samples into the Agilent 4200 TapeStation instrument. Carefully remove caps of tube strips.
6.13.12. Place ladder in position A1 on tube strip holder in the 4200 TapeStation instrument.
6.13.13. Select required sample positions on the 4200 TapeStation Controller Software.
6.13.14. Click Start.
6.13.15. The Agilent Tapestation Analysis Software opens after the run and displays results.
6.13.16. Use the estimated Average size of the library to calculate the nM concentration of library.

7. **Library dilution and Pooling** (Proceed to this step only if starting the run on same day)
   7.1. Fill the two Qubit concentration values and average library size estimated from TapeStation to the “Normalization & Multiplexing” Tab of worksheet. Dilute all the libraries according to the “Normalization & Multiplexing” tab in worksheet calculation to 10 nM.
   7.2. Once all the libraries diluted to 10 nM, pool the libraries to a single tube as per the specified volume in the worksheet for each sample. Pooling will be based on how much data required to generate per sample. If all the libraries are of same capture size, pool equal volume of libraries.
   7.3. Prepare at least 150µL of pooled 10nM library
   7.4. Proceed to sequencing run set up.
