## Supplemental Table 1 for "SARS-CoV-2 Whole Genome Amplification and Sequencing for Effective Population-Based Surveillance and Control of Viral Transmission"

**Supplemental Table 1.** M13-Tagged PCR Primers used for entire SARS-CoV-2 genome amplification*.

**Primer Name	Sequence (5’-3’)	Region	Size (bp)**

| WHCV_M13-F1 | GTAAAACGACGGCCAGTCCAGGTAACAAACCAACCAACTT | 36-58 | 1494 |
| --- | --- | --- | --- |
| WHCV_M13-R1 | CAGGAAACAGCTATGACCGGCAACCAACATAAGAGAACACAC | 1507-1530 |  |
| WHCV_M13-F2 | GTAAAACGACGGCCAGTCAACCAAATGTGCCTTTCAACTC | 1217-1239 | 1554 |
| WHCV_M13-R2 | CAGGAAACAGCTATGACCCACAGTGTCATCACCAAAAGTAACCT | 2746-2771 |  |
| WHCV_M13-F3 | GTAAAACGACGGCCAGTTGTCACGCACTCAAAGGGATT | 2408-2428 | 1404 |
| WHCV_M13-R3 | CAGGAAACAGCTATGACCGACAGCTAAGTAGACATTTGTGCGAA | 3787-3812 |  |
| WHCV_M13-F4 | GTAAAACGACGGCCAGTATGCCATGCAAGTTGAATCTGAT | 3523-3545 | 1503 |
| WHCV_M13-R4 | CAGGAAACAGCTATGACCTGCGTGTGGAGGTTAATGTTGT | 5005-5026 |  |
| WHCV_M13-F5 | GTAAAACGACGGCCAGTGATCTCTCAAAGTGCCAGCTACAGT | 4681-4705 | 1518 |
| WHCV_M13-R5 | CAGGAAACAGCTATGACCTTATAATCAATAGCCACCACATCACC | 6174-6199 |  |
| WHCV_M13-F6 | GTAAAACGACGGCCAGTAGAAACTTTGTATTGCATAGACGGTG | 5807-5832 | 1269 |
| WHCV_M13-R6 | CAGGAAACAGCTATGACCACCAGTACAGTAAGAAGGCATGCC | 7053-7076 |  |
| WHCV_M13-F7 | GTAAAACGACGGCCAGTGTTTAGCTGCTGTTAATAGTGTCCCTT | 6658-6684 | 1384 |
| WHCV_M13-R7 | CAGGAAACAGCTATGACCTGCAACTTCCGCACTATCACC | 8022-8042 |  |
| WHCV_M13-F8 | GTAAAACGACGGCCAGTTCCTACTGACCAGTCTTCTTACATCGT | 7727-7753 | 1525 |
| WHCV_M13-R8 | CAGGAAACAGCTATGACCTTTCACAAGTGCCGTGCCTAC | 9232-9252 |  |
| WHCV_M13-F9 | GTAAAACGACGGCCAGTGGTTTGCCTGGCACGATATTAC | 8883-8904 | 1488 |
| WHCV_M13-R9 | CAGGAAACAGCTATGACCACTTAGGTGTCTTAGGATTGGCTGTAT | 10345-10371 |  |
| WHCV_M13-F10 | GTAAAACGACGGCCAGTTTGTCATCTCGCAAAGGCTCT | 9974-9994 | 1523 |
| WHCV_M13-R10 | CAGGAAACAGCTATGACCGAGATTATAAGAGCCCACATGGAAA | 11473-11497 |  |
| WHCV_M13-F11 | GTAAAACGACGGCCAGTGCTATGGGTATTATTGCTATGTCTGCT | 11124-11150 | 1453 |
| WHCV_M13-R11 | CAGGAAACAGCTATGACCTGGATTTCCCACAATGCTGAT | 12557-12577 |  |
| WHCV_M13-F12 | GTAAAACGACGGCCAGTCTGATCAAGCTATGACCCAAATGT | 12295-12318 | 1451 |
| WHCV_M13-R12 | CAGGAAACAGCTATGACCGCAACAGCTGGACAATCCTTAAGT | 13723-13746 |  |
| WHCV_M13-F13 | GTAAAACGACGGCCAGTTCTGCGGTATGTGGAAAGGTTAT | 13396-13418 | 1191 |
| WHCV_M13-R13 | CAGGAAACAGCTATGACCGTCAGCAGCATACACAAGTAATTCCT | 14562-14587 |  |
| WHCV_M13-F14 | GTAAAACGACGGCCAGTAGGGCTTTAACTGCAGAGTCACAT | 14201-14224 | 1418 |
| WHCV_M13-R14 | CAGGAAACAGCTATGACCGCGGACATACTTATCGGCAATT | 15598-15619 |  |
| WHCV_M13-F15 | GTAAAACGACGGCCAGTTCAATAGCCGCCACTAGAGGAG | 15188-15209 | 1420 |
| WHCV_M13-R15 | CAGGAAACAGCTATGACCTCACCAGCATTTGTCCAGTCAC | 16587-16608 |  |
| WHCV_M13-F16 | GTAAAACGACGGCCAGTTTGGGGCTTGTGTTCTTTGC | 16257-16276 | 1512 |
| WHCV_M13-R16 | CAGGAAACAGCTATGACCCAAGCAGGGTTACGTGTAAGGAAT | 17746-17769 |  |
| WHCV_M13-F17 | GTAAAACGACGGCCAGTTGTCAATGCCAGATTACGTGCT | 17410-17431 | 1507 |
| WHCV_M13-R17 | CAGGAAACAGCTATGACCTAACAAAGCACTCGTGGACAGC | 18896-18917 |  |
| WHCV_M13-F18 | GTAAAACGACGGCCAGTTATGGGCACATGGCTTTGAGT | 18609-18629 | 1454 |
| WHCV_M13-R18 | CAGGAAACAGCTATGACCTAAGAACACCATTACGGGCATTT | 20041-20063 |  |
| WHCV_M13-F19 | GTAAAACGACGGCCAGTTTGATGGACAACAGGGTGAAGTAC | 19680-19703 | 1556 |
| WHCV_M13-R19 | CAGGAAACAGCTATGACCCGAAGTGTCCCATGAGCTTATAAA | 21213-21236 |  |
| WHCV_M13-F20 | GTAAAACGACGGCCAGTAGGAGTTGCACCAGGTACAGCT | 20902-20923 | 1480 |
| WHCV_M13-R20 | CAGGAAACAGCTATGACCACCCACATAATAAGCTGCAGCAC | 22360-22382 |  |
| WHCV_M13-F21 | GTAAAACGACGGCCAGTCTATTAATTTAGTGCGTGATCTCCCTC | 22204-22230 | 1527 |
| WHCV_M13-R21 | CAGGAAACAGCTATGACCAAATTTGTGGGTATGGCAATAGAGTTA | 23705-23731 |  |
| WHCV_M13-F22 | GTAAAACGACGGCCAGTACTTACTCCTACTTGGCGTGTTTATTC | 23462-23488 | 1637 |
| WHCV_M13-R22 | CAGGAAACAGCTATGACCGCATTAATGCCAGAGATGTCACC | 25077-25099 |  |
| WHCV_M13-F23 | GTAAAACGACGGCCAGTCTATCATCTTATGTCCTTCCCTCAGTC | 24716-24742 | 1503 |
| WHCV_M13-R23 | CAGGAAACAGCTATGACCTAGTCGTCGTCGGTTCATCATAAAT | 26195-26219 |  |
| WHCV_M13-F24 | GTAAAACGACGGCCAGTTACTTCAGGTGATGGCACAACAA | 25915-25937 | 1545 |
| WHCV_M13-R24 | CAGGAAACAGCTATGACCAAGCTCACAAGTAGCGAGTGTTATCA | 27435-27460 |  |
| WHCV_M13-F25 | GTAAAACGACGGCCAGTCGTGTAGCAGGTGACTCAGGTTT | 27094-27116 | 1493 |
| WHCV_M13-R25 | CAGGAAACAGCTATGACCTACCGTCACCACCACGAATTC | 28567-28587 |  |
| WHCV_M13-F26 | GTAAAACGACGGCCAGTGGACCCCAAAATCAGCGAAAT | 28302-28322 | 1562 |
| WHCV_M13-R26 | CAGGAAACAGCTATGACCAAAATCACATGGGGATAGCACTACT | 29840-29864 |  |

*Primers were modified from Wu et al 2020 by adding M13 tails.
